# Loss of generalist plant species and functional diversity decreases the robustness of a seed dispersal network

**DOI:** 10.1101/187179

**Authors:** Vinicius A. G. Bastazini, Vanderlei J. Debastiani, Bethânia O. Azambuja, Paulo R. Guimarães, Valério D. Pillar

## Abstract

Understanding cascading effects of species loss has become a major challenge for ecologists. Traditionally, the robustness of ecological networks has been evaluated based on simulation studies where primary extinctions occur at random or as a function of species specialization, ignoring other important biological factors. Here, we estimate the robustness of a seed dispersal network from a grassland–forest mosaic in southern Brazil, simulating distinct scenarios of woody plant species extinction, including scenarios where species are eliminated based on their evolutionary and functional distinctiveness. Our results suggest that the network is more robust when species are eliminated based on their evolutionary uniqueness, followed by random extinctions, the extinction of the most specialist species, functional distinctiveness and, at last, when the most generalist species are sequentially eliminated. Our results provide important information for grassland–forest mosaic management, as they indicate that loss of generalist species and functional diversity makes the system more likely to collapse.

## Introduction

As we face the prospect of an unprecedented anthropogenic mass extinction, with species being lost at rates that are two or three orders of magnitude greater than the background rates from the geological record (Pimm et al. 2014, Ceballos et al. 2015), understanding and predicting the consequences of species extinction has become an urgent task for conservation scientists and practitioners (Vieira & Almeida-Neto 2014, Ceballos et al. 2015). Most empirical studies examining the magnitude of biodiversity loss have usually ignored coextinction scenarios (Dunn et al. 2009, Vieira & Almeida-Neto 2014), where the loss of one or more species may unleash a cascade of secondary extinctions, that may bring entire communities and ecosystems to collapse (Jackson et al. 2001, Fowler 2010, Colwell et al. 2012, Säterberg et al. 2013, Brodie et al. 2014, Donoso et al. 2017). Although the magnitude of coextinction process is still virtually unknown, given the difficulties of documenting it in natural systems (see for instance Dunn et al. 2009 and Moir et al. 2010), it has been suggested that these cascading effects should be more pervasive in some types of ecological interactions, such as parasitism and mutualism (Dunn et al. 2009, Kiers et al. 2010). The recent and rapid rise of Network Ecology – i.e., a field of research interested in understanding the function, structure, and evolution of ecological systems, using network models and analyses (Borrett et al. 2014) – has proved itself to be a powerful tool to describe the complexity of species interactions and their interdependence (Proulx et al. 2005, Pascual and Dunne 2006, Miranda et al. 2013), including their response to disturbance and cascading effects, such as coextinctions. Understanding what determines the robustness of mutualistic networks, i.e., the system’s tolerance to secondary extinctions, is currently one of the main challenges faced by network ecologists (Solé & Montoya 2001, Dunne et al. 2002, Memmot et al. 2004, Burgos et al. 2007, Rezende et al. 2007, Mello et al. 2011a, b, Pocock et al. 2012, Eklöf et al. 2013, Vieira et al. 2013, Astegiano et al. 2015). However, despite their importance, most studies trying to estimate the robustness of ecological networks have been traditionally based on scenarios that ignore other important ecological and evolutionary factors, such as species traits, associated to the likelihood of a species becoming extinct (e.g., Dunne et al. 2002, Memmot et al. 2004, Burgos et al. 2007, Pocock et al. 2012, but see Curtsdotter et al. 2011, Srinivasan et al. 2007, Vieira et al. 2013, Astegiano et al. 2015). In these studies, primary species extinctions are assumed to occur at random or as a consequence of species specialization (Solé & Montoya 2001, Dunne et al. 2002, Memmot et al. 2004, Burgos et al. 2007, Rezende et al. 2007, Mello et al. 2011a, b, Pocock et al. 2012).

Species functional traits – behavioral, morphological or ecophysiological characteristics associated with biotic interactions, environmental conditions and/or ecosystem functions (Schmitz et al. 2015, Lefcheck et al. 2015) – play an important role in determining the structure and stability of ecological networks, as species traits can directly constrain the likelihood of interactions among individuals or species (Santamaría & Rodríguez-Gironés 2007, Vizentin-Bugoni et al. 2014, Bastazini 2017) and the probability of extinction of a species (Purvis et al. 2000, Cardillo et al. 2005, 2008, Reynolds et al. 2005). Species traits imply that species are different in their ecological requirements and their effects on ecosystems. However, they are not equally different (Lefecheck et al. 2015), and some species are more likely to retain similar or “redundant” traits (Rosenfeld 2002, Pillar et al. 2013, Kang et al. 2015), conferring higher resistance and resilience to the system, in what is know as the “insurance effect of biodiversity” (Walker 1992, Pillar et al. 2013, Kang et al. 2015, Oliver et al. 2017). Thus, one might expect that the loss of more “distinct” and less redundant species in term of their traits might have a greater effect on how ecological networks respond to disturbances, as their role in the network can not be compensated by other species.

Plant–animal seed disperser interactions are an important mutualism that shapes both, ecological and evolutionary dynamics, through demographic processes that affect the fitness, reproduction success and the distribution of partner species (Snow 1971, Morton 1973, Herrera 1985, Jordano 2000, Correa et al. 2015). That these interactions are key drivers of biodiversity patterns is supported by the fact that only in the Tropics, up to 90% of all plant species rely on animals to disperse their seeds (Jordano 2000). Given their significance to maintain biodiversity and support ecosystem functioning, plant–disperser interactions are considered a crucial component in conservation and restoration strategies (Wunderle Jr. et al. 1997, Trakhtenbrot et al. 2005). Insofar, few studies have evaluated the robustness of seed dispersal systems (but see Mello et al. 2011a, b, Timóteo et al. 2016, Costa et al. 2018).

Seed dispersal networks also play a critical role in terrestrial ecotone zones, such as in grassland-forest ecotones, as animal seed disperser may help plant species to expand their distribution across phytophysiognomy boundaries, ultimately transforming the landscape (Jensen et al. 1986, Bossuyt et al. 1999, Fragoso et al. 2003, Azambuja 2009, Carlucci et al. 2011, Müller et al. 2012, Myster 2012). Grasslands are one of the major biomes of the planet, occupying an area equivalent to 31-43% of the land surface (Coupland 1979, White et al. 2000). Grasslands support high levels of biological diversity and endemism (White et al. 2000, Bond & Parr 2010, Iganci et al. 2011, Parr et al. 2014) and are responsible for maintaining important ecosystem processes and economic activities, such as acting as carbon sinks, supporting livestock production and providing aesthetic and cultural values to local communities, especially pastoral societies (Coupland 1979, Breymer & Van Dyne 1980, White et al. 2000, Hu et al. 2001, Overbeck et al. 2007, Curtin & Western 2008, Parr et al. 2014, Henderson et al. 2016). Nonetheless, their conservation has been historically neglected by environmental policies (Overbeck et al. 2007, Bond & Parr 2010, Parr et al. 2014) and despite their large area, and economic and environmental value, grasslands are among the most altered and threatened ecosystems of the planet (White et al. 2000, Bond & Parr 2010, Henwood et al. 2010, Parr et al. 2014). In addition to anthropogenic threats, such as overgrazing, farm intensification and afforestation, temperate grasslands and savannas suffer from a natural process of forest expansion (Schwartz et al. 1996, Bowman et al. 2001, Bond & Parr 2010, Overbeck et al. 2007, Müller et al. 2012). Although forest expansion is considered a threat to grassland conservation, this process creates a unique and useful scenario for ecological studies, as grassland–forest ecotones encompass a “mosaic of life forms” that share very distinct evolutionary origins and life histories (Luza et al. 2015) that compete for space and other resources. Seed dispersal networks, formed by woody plants and frugivores, are likely to be an important driver of spatial and temporal dynamics in grassland–forest mosaics, as seed dispersal of woody plants may accelerate the expansion of forest ecosystems (Azambuja 2009, Carlucci et al. 2011, Müller et al. 2012, Myster 2012). Thus, grassland–forest ecotones make an interesting case scenario to study simulated coextinctions, with important consequences to the development of conservation policies. For instance, if the goal of conservation efforts is to protect grasslands, these analyses can help us to identify species that should be managed in order to maintain grasslands. Also, if conservation efforts are aimed to maintain forest expansion or forest species, this sort of analyses can help us to identify species that should be of higher priority in conservation.

Here, we estimated the robustness of a seed dispersal network formed by woody plants and birds from a grassland–forest mosaic from southern Brazil. We simulated five distinct scenarios of woody plant species extinction, including scenarios where species are eliminated based on their evolutionary and functional distinctiveness. Our main hypothesis is that the loss of more functional distinct, and consequently less redundant, species should have a large effect on network robustness, as their role in the network can not be compensated by the remaining species. Given the lack of phylogenetic signal in traits in this network (Bastazini et al. 2017), we also expect that loss of phylogenetic distinct species will be less important. Although the role of bottom-up and top-down ecosystem regulation has been a much-debated topic for decades now (see for instance Hanley & La Pierre 2015 and references therein), both theoretical and empirical evidence suggest that bottom-up dynamics are more prominent in shaping community dynamics in multitrophic scenarios (Goudard & Loreau 2008, Scherber et al. 2010). Moreover, recent evidence suggests that plant extinctions may be more likely to trigger animal coextinctions than vice versa (Schleuning et al. 2016). Thus, we assumed a bottom-up regulation and simulated distinct scenarios of primary woody plant extinction and their effects on bird coextinctions.

## Material and Methods

### Study system and seed dispersal network

Our study system comprises a seed dispersal network from a forest–grassland mosaic located in southern Brazil (Azambuja 2009, Bastazini et al. 2017). The site (between 30°25’03”S 52°21’37”W and 30°25’54”S 52°22’40”W) is characterized by gently rolling terrains, with altitudes ranging from 100 to 210m, and by a grassland matrix, which is manly used for cattle ranching (Azambuja 2009). Large forest patches occupy hill slopes and riparian areas and smaller shrubby and forest patches are sparsely distributed across the grassy landscape (Azambuja 2009, Bastazini et al. 2017). The average annual temperature is 16.5°C and the mean annual precipitation is 1504 mm (Instituto de Pesquisas Agronômicas 1989). For more details on the study region, please refer to Azambuja (2009).

We used data collected during a bird trapping study that lasted one year, between July 2007 and June 2008 (Azambuja 2009, Bastazini et al. 2017). Birds were captured in mist nets, which were set up for eight consecutive days every month in grasslands, approximately 1 m away from the border of forest patches, in order to optimize bird capture success. Captured birds were placed into fabric bags for 20 minutes, so their feces could be collected (Azambuja 2009, Bastazini et al. 2017). The seeds found in fecal samples were identified to the finest taxonomic resolution possible. Based on this information, we built a qualitative interaction matrix comprising of 22 woody plant species and 12 frugivorous bird species (Fig. 1). In a recent contribution, Bastazini et al. (2017) demonstrated that trait coupling between birds and plants is an important driver of the structure of this network. We used the phylogeny generated by Bastazini et al. (2017), based on the APG III megatree (R20100701.new; Angiosperm Phylogeny Group 2009), and the traits used in their analyses, two continuous (diaspore diameter and maximum plant height) and three categorical traits (aril presence, diaspore shape and color), to estimate the phylogenetic and functional distinctiveness index explained in detail below. Further information on the phylogeny and the functional traits used in our analyses can be found in Bastazini et al. (2017) and their Supplementary Material 4.

**Fig. 1.**
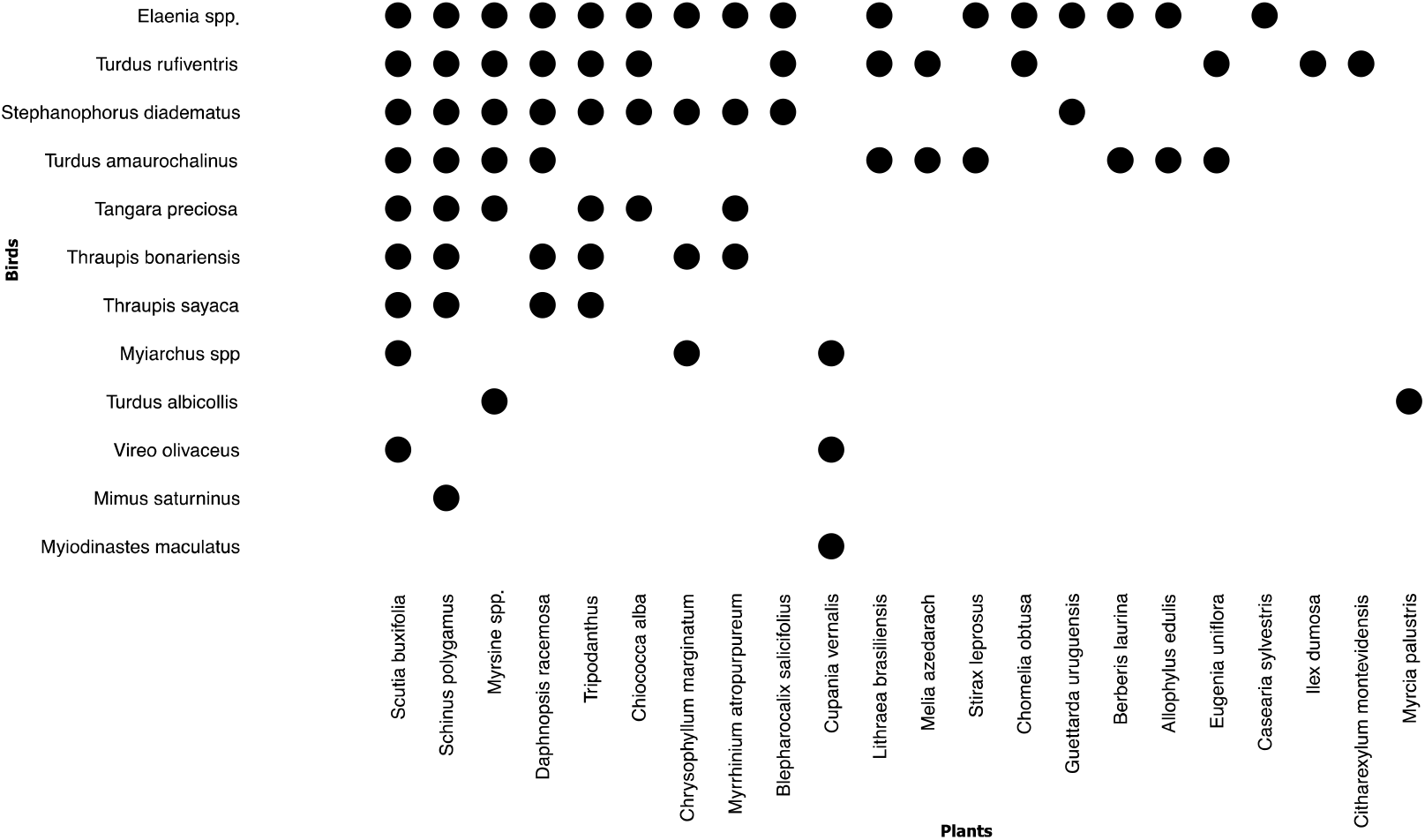
Bi-adjacency matrix representing the seed dispersal network from a forest-grassland mosaic in southern Brazil.

### Numerical analyses

We estimated the phylogenetic distinctiveness of each plant species, i.e., how isolated a species is on the phylogenetic tree, using the Fair Proportion metric, which is defined as the sum of all edge lengths between the species and the root of the phylogenetic tree, with each edge length being divided by the number of species in the cluster it subtends (Redding et al. 2008). We repeated the same procedure in order to estimate the functional distinctiveness of each species, i.e., how distinct a species is in terms of its functional traits, based on a functional dendrogram using plant traits. The functional dendrogram was built using Ward’s sum of squares clustering procedure based on an Euclidean distance matrix calculated from species trait values (Legendre & Legendre 2012).

Secondary extinctions of woody plants were simulated by five distinct elimination schemes. The first was based on **i)** random extinctions (using 1,000 simulations) and provided a baseline scenario to compare the effects of the other scenarios (Dunne et al. 2002, Memmot et al. 2004, Astegiano et al. 2015). The other scenarios considered species specialization in the seed dispersal network, either with **ii)** the most generalist (most connected plants) or **iii)** the most specialist (poorly connected plants) species disappearing first; iv) species eliminated based on their evolutionary distinctiveness; and **v)** based on species’ functional distinctiveness. In both, **iv)** and **v)**, most distinct species, either in terms of their traits of phylogenetic position are eliminated first. For each scenario we calculated network robustness (*R*) defined as the area below the Attack Tolerance Curve (ATC; Albert & Barabási 2002, Burgos et al. 2007), which represents the curve of the fraction of surviving bird species as a function of the eliminated plant species. *R* values vary between 0 and 1, where values closer to 1 indicate higher network robustness (Burgos et al. 2007).

All numerical simulations and analyses were performed in the R environment (R Core Team 2012) and the code is available on Github (https://github.com/bastazini/Networks).

## Results

Species were more variable in terms of their phylogenetic distinctiveness (range: 0.19 – 0.48) than in terms of their functional distinctiveness (range: 0.19 – 0.25; Fig. 2). There was no correlation between both measures of distinctiveness (Pearson’s r = −0.015; P-value = 0.94), which indicates that most unique species in the phylogenetic tree are not the most unique species in terms of their functional traits (Fig. 2).

**Fig. 2.**
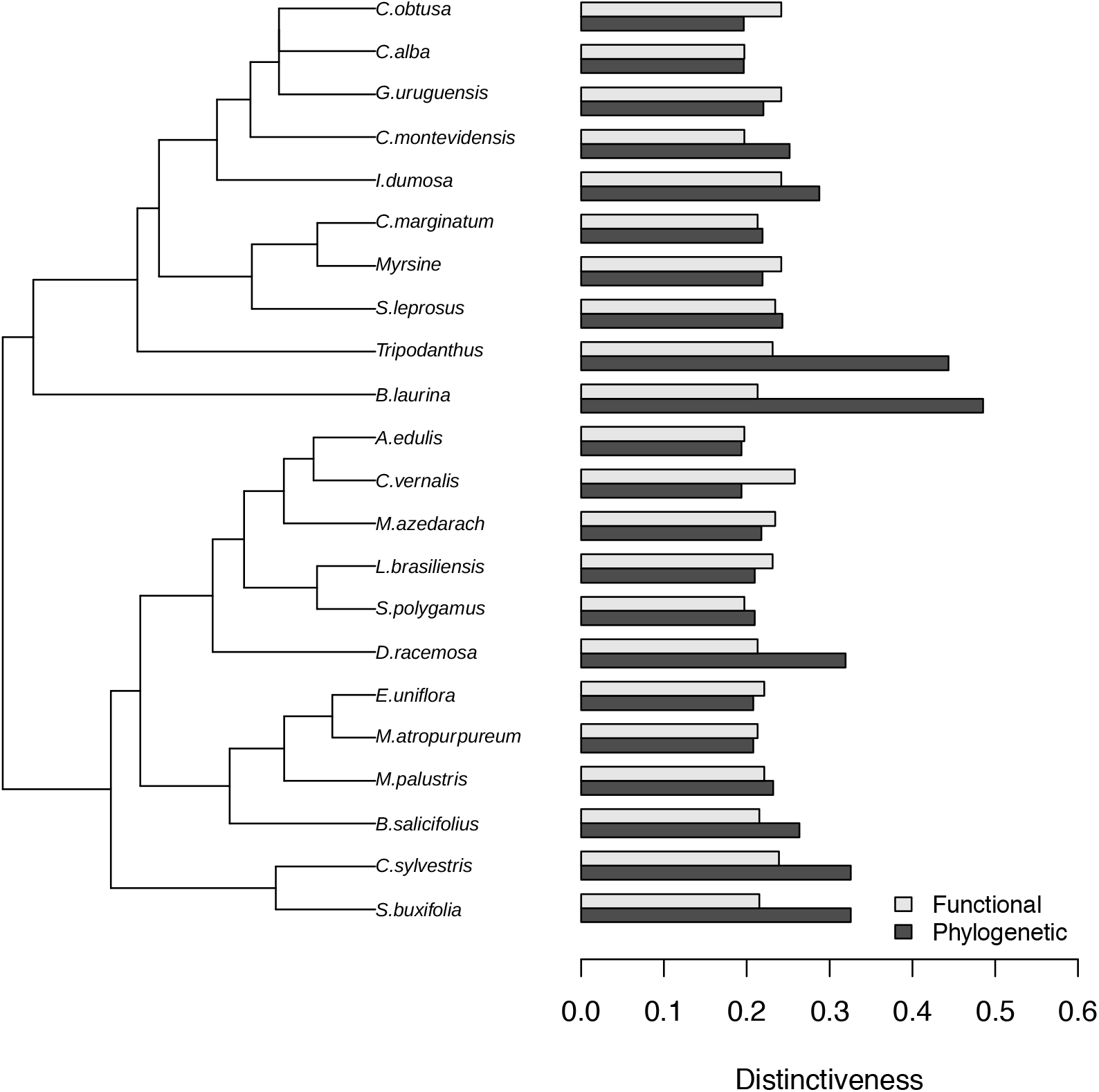
Functional and phylogenetic distinctiveness of plant species, with the phylogeny plotted alongside, in a seed dispersal network from southern Brazil.

Considering all five scenarios, our seed dispersal network was likely to be very robust to the loss of woody plant species (average robustness considering all five simulated scenarios = 0.74 ± SD = 11.85). However, our simulations showed that distinct extinction scenarios have distinct effects on network robustness (Fig. 3). The loss of generalist species was more detrimental to network robustness, as the ATC of this scenario exhibited a very abrupt decline (Fig. 3 ii). The other scenarios showed a less steep response to the loss of plant species. Network robustness was higher when species were eliminated based on their evolutionary uniqueness, followed by random extinctions, the extinction of the most specialist species and functional distinctiveness (Fig. 3).

**Fig. 3.**
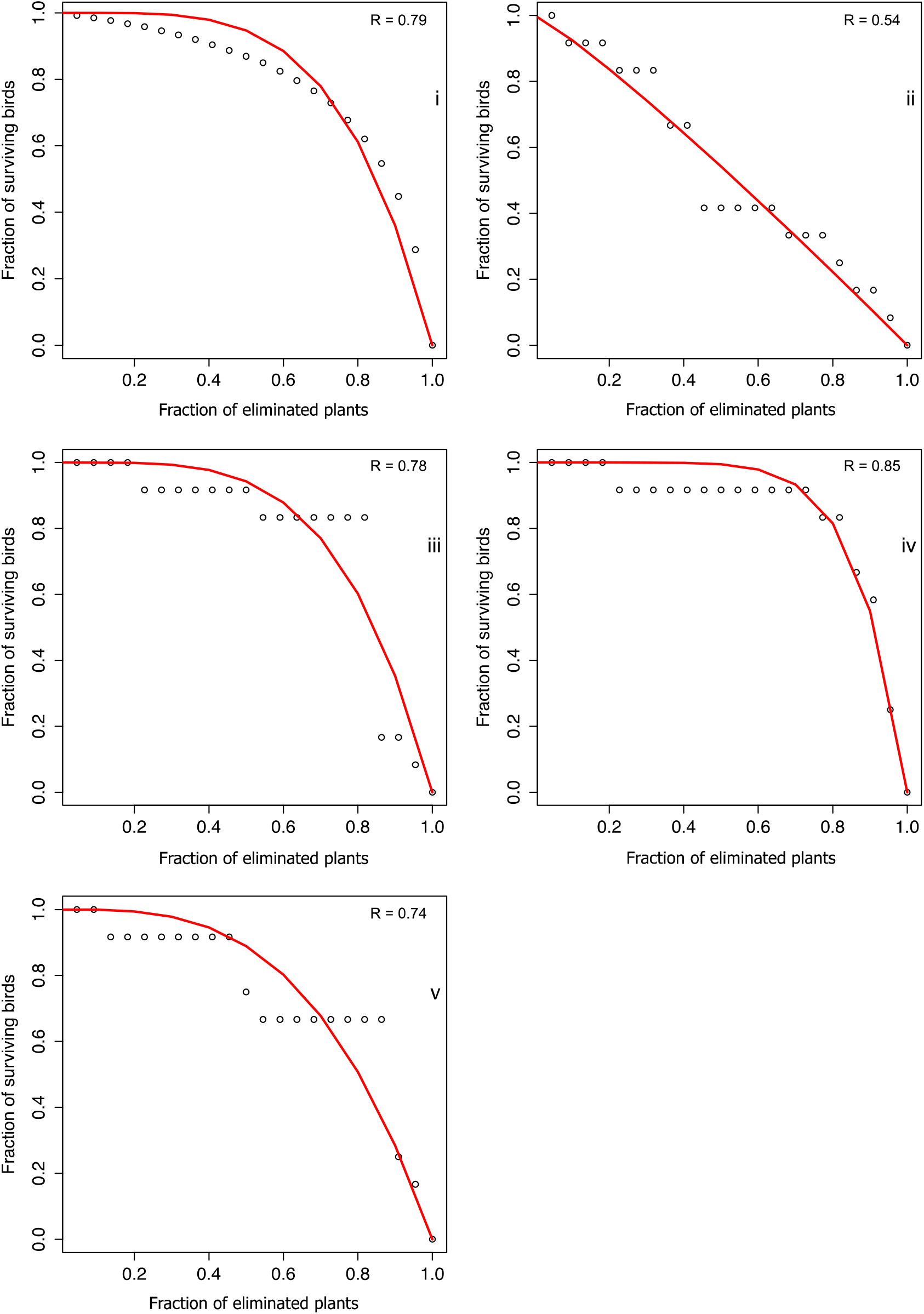
Attack Tolerance Curves under different plant extinction sequences: i) random; extinctions based on species specialization either with ii) the most generalist (most connected plants) or iii) the most specialist (poorly connected plants) species disappearing first; iv) species evolutionary distinctiveness; v) species functional distinctiveness.

## Discussion

Over the past years, there has been a growing interest in trying to understand and estimate the consequences of coextinctions in ecological networks (e.g., Solé & Montoya 2001, Dunne et al. 2002, Memmot et al. 2004, Burgos et al. 2007, Rezende et al. 2007, Mello et al. 2011a, b, Pocock et al. 2012). Insofar, most empirical evaluations have been based on scenarios where primary extinctions occur as a function of species specialization or as random events (Solé & Montoya 2001, Dunne et al. 2002, Memmot et al. 2004, Burgos et al. 2007, Pocock et al 2012, Vieira et al. 2013). These scenarios may be unrealistic or offer a partial understanding of coextinction processes, as other ecological and evolutionary factors, such as body size and geographical range, influence the probability of a species becoming extinct (Purvis et al. 2000, Cardillo et al. 2005, 2008, Reynolds et al. 2005). Here, we provide further developments to coextinction analyses based on ATCs, combining functional and phylogenetic information, and show that the loss of generalist species has a larger effect on network robustness when compared to other extinction scenarios. Vieira et al. (2013) investigated the functional and phylogenetic consequences of random pollinator extinction in several pollination networks using a different metric to quantify functional and phylogenetic uniqueness. Their results suggest that there is an uncoupled response between functional and phylogenetic loss in mutualistic systems. Our study corroborates their findings, since our extinction scenarios based on functional and phylogenetic distinctiveness shows very different responses in terms of network robustness. As we predicted, given the lack of phylogenetic signal in plant traits in this seed dispersal network (Bastazini et al. 2017), plant functional and phylogenetic effects on bird species extinction were uncoupled, with the loss of phylogenetic distinct plant species having a lower impact on network robustness.

Despite this growing interest in coextinction and robustness analyses, studies focusing on seed dispersal networks are still scarce. Mello et al. (2011b) analyzed the robustness of 21 seed dispersal networks by simulating random extinctions of plants and frugivores (bats and birds). For plant–bird networks, they found that networks should be very robust to the loss of plant species, with a mean value of robustness (*R* = 0.75) similar to our result. Future studies are necessary to demonstrate whether seed dispersal networks are as robust as demonstrated by Mello et al. (2011b) and our own analyses. But as plant-seed disperser interactions are a system of low specialization (Herrera 1995, Muller-Landau & Hardesty 2005, Donoso et al. 2017), high network robustness should be expected. The sensitiveness of dispersal networks to the primary extinction of plants or seed dispersers is still unclear. However Mello et al. (2011b) suggested that networks should be less robust when animal species are lost, Schleuning et al. (2016) argued that plant extinctions are more detrimental than animal extinctions in mutualistic networks. An important further step is to investigate the ability of species to replace their lost mutualistic partner. Experimental studies have suggested that rewiring, i.e., the re-arrangement of species interactions through time, is likely to promote higher resistance of seed dispersal networks (Timóteo et al. 2016, Costa et al. 2018). Nonetheless, we still lack a deep understanding of the underlying mechanisms driving rewiring in mutualistic networks, as different factors such as spatial-temporal cooccurrence, environmental gradients and species traits and abundance may determine the probability of species interactions (Danielson 1991, Jordano 2000, Vizentin-Bugoni et al. 2014, Schmitz et al. 2015, Bastazini et al. 2017). Although network rewiring models are likely to lead to more accurate predictions (Timóteo et al. 2016, Costa et al. 2018), not being able to correctly account for the mechanisms that determine the re-arrangement of species interactions or simulating unconstrained rewiring might lead to an overestimation of network robustness to secondary extinctions (Costa et al. 2018), which is undesirable from a conservation stand point. Nonetheless, we stress that although understanding the role of network rewiring is a critical question yet to be answered, our static and conservative approach provides valuable insights on the effects of eco-evolutionary mechanisms on network disassembly.

It is well recognized that generalist species, i.e., species with larger number of connections, play an important role in ecosystem functioning and stability (Memmot et al. 2004, Richmond et al. 2005, Gaston & Fuller 2008, Poisot et al. 2013, Valiente-Banuet et al. 2015). González et al. (2010) suggested that generalist species have, at least, two important roles in pollination networks; the first obvious and intuitive role is that generalist species are able to interact with more species than specialist ones and the second role is the ability to connect otherwise unconnected sub-networks. Together, these two roles imply that generalist species are responsible for creating a more cohesive network. This cohesive pattern is likely to increment the ability of the system to respond to perturbations, and consequently increase its stability (Bascompte et al. 2003). Generalist and common species should receive special attention in conservation as even small declines in their populations may result in significant disruption of ecosystems (Gaston & Fuller 2008). Our seed dispersal network comprises a very heterogeneous set of plant species in terms of phylogenetic history (15 taxonomic families), life history and functional groups, including trees (e.g., *Scutia buxifolia, Cupania vernalis* and *Styrax leprosus*), shrubs *(Berberis laurina, Chomelia obtusa* and *Daphnopsis racemosa*), vines (*Chiococca* alba) and hemiparasite species (*Tripodanthus* spp.). Among this diverse set of taxa, five of them are responsible for approximately 50% of all pairwise interactions, showing a larger number of connections: *Scutia buxifolia, Schinus polygamus, Myrsine* spp., *Daphnopsis racemosa* and *Tripodanthus* spp. Overall, these five taxa are characterized by low functional distinctiveness, with values lower than the median (except for *Tripodanthus* spp. and *Myrsine* spp.) and, except for *Schinus polygamus*, high phylogenetic distinctiveness (with values above the median). A common trait shared by these species is the size of their fruits, which is relatively small compared to the other species. As in other trophic interactions where predators have to swallow whole food items, fruit size is an important constraint in seed dispersal networks, and as a general rule, plants with large seeds and fruits tend to attract fewer animal species than plants with small fruits (Wheelwright 1985, Jordano 2000, Dehling et al. 2016, Donoso et al. 2017). Temporal availability and co-ocurrence with its seed dispersers have also been found to be important factors that increment the chance of a plant being consumed and dispersed by an animal (Jordano 2000). Among these five taxa, *Myrsine* spp. is the only one that produces fruits almost year-round (Azambuja 2009, Bastazini et al. 2017). Also, these well-connected species are among the taxa with larger temporal co-occurrence with the most bird species (Bastazini et al. 2017). Thus, small fruits and phenological patterns seemed to be a possible explanation to some structural patterns and the robustness of this seed dispersal network.

At last, we would like to underline the implications of our study to conservation policies, especially in southern Brazil. The conservation of grassland–forest mosaics of southern Brazil has been the focus of much debate in recent years (Overbeck et al., 2007, 2013, Luza et al. 2014). As in other grassland ecosystems around the World, the grasslands of southern Brazil have experienced an increase in density of woody species that is drastically changing its physiognomy (Overbeck et al 2007, Müller et al. 2012). Forest expansion seems to be influenced by nucleation processes influenced by the presence of facilitating structures, such as rocky outcrops or isolated trees established in the grassland matrix, that act as perches for vertebrate dispersers and as safe sites for woody plants (Carlucci et al., 2011, Müller et al. 2012). Most of the generalist tree species that we found in our study are important species for woody plant encroachment of grasslands of southern Brazil (Carlucci et al. 2011, Müller et al. 2012). As we found the loss of generalist species and the loss of functional uniqueness are more detrimental to the robustness of seed dispersal networks of southern Brazil, these results could help to develop sounder strategies in species management and restoration of grassland–forest mosaics in southern Brazil.

In summary, our findings provide important information for forest and grassland management in southern Brazil, as they indicated that the sequential extinction of generalist woody plant species and functional plant diversity made the system more likely to collapse, whereas the loss of the most “unique” species, in terms of their evolutionary history, and specialist species had a smaller effect on network robustness. Moreover, despite its simplicity, our framework stresses the importance of considering distinct extinction scenarios and can help ecologists to understand and predict cascading effects in ecological systems.

## Acknowledgments

We thank Andreas Kindel, André L. Luza, Danis Kiziridis, Kirsten Henderson, Paulo I. K. L. Prado, Rodrigo S. Bergamin and Sandra C. Muller and two anonymous reviewers for revising and providing valuable suggestions that greatly improved this manuscript. VAGB received support from CAPES (grant #1002302), the TULIP Laboratory of Excellence (ANR-10-LABX-41 and 394 ANR-11-IDEX-002-02) and from a Region Midi-Pyrenees project (CNRS 121090). VP received support from CNPq, Brazil (grant # 307689/2014-0). PRG was supported by FAPESP (grant #2009/54422-8) and CNPq.

